# Direct interaction of PIWI and DEPS-1 is essential for piRNA function and condensate ultrastructure in *Caenorhabditis elegans*

**DOI:** 10.1101/580043

**Authors:** KM Suen, F Braukmann, R Butler, D Bensaddek, A Akay, C. -C Lin, N Doshi, A Sapetschnig, A Lamond, JE Ladbury, EA Miska

**Affiliations:** Wellcome Trust Cancer Research UK Gurdon Institute, University of Cambridge, Tennis Court Road, Cambridge CB2 1QN, UK; Department of Genetics, University of Cambridge, Downing Street, Cambridge, CB2 3EH, UK; Wellcome Sanger Institute, Wellcome Trust Genome Campus, Hinxton, CB10 1SA, UK; Laboratory for Quantitative Proteomics, Centre for Gene Regulation and Expression, College of Life Sciences, University of Dundee, Dow Street, Dundee DD1 5EH, UK; School of Molecular and Cellular Biology, University of Leeds, LC Miall Building, Leeds, LS2 9JT, UK; University of Cambridge, School of Clinical Medicine, Cambridge Biomedical Campus, Cambridge, CB2 0SP, UK; Bioscience Core labs, King Abdullah University of Science and Technology, Thuwal 23955-6900 Saudi Arabia

## Abstract

Membraneless organelles are platforms for many aspects of RNA biology including small non-coding RNA (ncRNA) mediated gene silencing. How small ncRNAs utilise phase separated environments for their function is unclear. To address this question, we investigated how the PIWI-interacting RNA (piRNA) pathway engages with the membraneless organelle P granule in *Caenorhabditis elegans*. Proteomic analysis of the PIWI protein PRG-1 revealed an interaction with the constitutive P granule protein DEPS-1. Furthermore we identified a novel motif on DEPS-1, PBS, which interacts directly with the Piwi domain of PRG-1. This protein complex forms intertwining ultrastructures to build elongated condensates *in vivo*. These sub-organelle ultrastructures depend on the Piwi-interacting motif of DEPS-1 and mediate piRNA function. Additionally, we identify a novel interactor of DEPS-1, EDG-1, which is required for DEPS-1 condensates to form correctly. We show that DEPS-1 is not required for piRNA biogenesis but piRNA function: *deps-1* mutants fail to produce the secondary endo-siRNAs required for the silencing of piRNA targets. Our study reveals how specific protein-protein interactions drive the spatial organisation and function of small RNA pathways within membraneless organelles.

## Introduction

The correct spatial organisation of molecules into organelles is essential for biological function. Recent studies revealed that membraneless organelles can be formed by proteins and nucleic acids condensing out of the bulk intracellular milieu, giving rise to liquid- or gel-like environments (Banani et al., 2017; Boeynaems et al., 2018; Shin and Brangwynne, 2017). Such phase separated organelles are platforms for different aspects of eukaryotic RNA biology: the nucleolus is required for the assembly of ribosomes (Feric et al., 2016), stress granules allow for translational stalling of mRNAs during stress-response (Molliex et al., 2015) and processing bodies (P-bodies) organise small RNA-mediated regulation of mRNA (Hubstenberger et al., 2017). However, the molecular mechanisms of how RNA and proteins in these phase separated organelles assemble remain largely unexplored.

Small non-coding RNAs (ncRNAs) execute diverse biological functions. mRNAs are targeted for silencing by small RNAs based on Watson-Crick base-pair complementarity in complex with members of the Argnonaute (Ago) protein family (Meister, 2013). Various Ago proteins associate with membraneless organelles such as the P-bodies and germ granules (Eulalio et al., 2007; Voronina et al., 2011). Indeed, recently it has been shown that human Ago2 together with its binding partner TNRC6B can form biomolecular condensates entirely on their own (Sheu-Gruttadauria and MacRae, 2018). A novel condensate, the Z granule, contains the Ago protein WAGO-4 to establish transgenerational inheritance (TEI) of RNAi in *C. elegans* (Ishidate et al., 2018; Wan et al., 2018). Hence, some small RNAs are routed through membraneless organelles. To further our understanding of how small RNAs operate within membraneless organelles, we turned to the piRNA pathway.

piRNAs associate with the PIWI clade proteins in the Argonaute family to repress transposable elements (TEs) (Czech et al., 2018; Malone and Hannon, 2009; Weick and Miska, 2014). Mutations in the piRNA pathway lead to varying degrees of infertility, indicating it plays an essential role in the survival of a species. For example, null mutations in each of the three *piwi*-coding genes lead to sterility in male mice (Carmell et al., 2007; Deng and Lin, 2002; Kuramochi-Miyagawa et al., 2004); depletion of the single functional PIWI protein in *C. elegans* leads to reduced fecundity (Simon et al., 2014); in humans, a mutation blocking the ubiquitination of the PIWI protein HIWI has been implicated in azoospermia (Gou et al., 2017).

Mature piRNAs mediate transcriptional and post-transcriptional gene silencing. In *C. elegans*, piRNAs are 21 nt long with a 5’ preference for U (Ruby et al., 2006) and contain 2-O-methylation at the 3’ end (Kamminga et al., 2012; Montgomery et al., 2012). piRNAs associates with the PIWI protein PRG-1 to scan for target mRNAs (Bagijn et al., 2013; Batista et al., 2008). Once discovered by the PRG-1/piRNAs complex, the target mRNAs serve as templates for the production of endo-siRNAs that are 22 nt long with a 5’ preference for G (22Gs) by the RNA-dependent-RNA polymerases (RdRPs) EGO-1 and RRF-1 (Das et al., 2008; Ashe et al., 2012; Bagijn et al., 2013; Shirayama et al., 2012). The *C. elegans* piRNA pathway offers a unique model for understanding how membraneless organelles engage with small RNA pathways as it requires two distinct and juxtaposed biomolecular condensates to achieve gene repression: the perinuclear-granules (P granules) where PRG-1 resides (Batista et al., 2008) and the secondary endo-siRNAs are entirely dependent on the mutator foci (Zhang et al., 2011; Phillips et al., 2012; Uebel et al., 2018).

Here we investigated how the piRNA pathway engages with the membraneless organelles P granules and mutator foci. Specifically, we determined the protein interactome of the *C. elegans* PIWI protein PRG-1. We found the P granule factor DEPS-1 directly binds to PRG-1 and identified a motif on DEPS-1 responsible for this interaction, which we termed the PIWI-binding site (PBS). Furthermore, we found that DEPS-1 and PRG-1 condensates are formed from smaller clusters of proteins that intertwine to produce elongated perinuclear granule. Removing the PBS from DEPS-1 leads to PRG-1 condensate to collapse into a compacted morphology. Surprisingly, mutating *deps-1* and *prg-*1 gives rise to aberrant mutator foci and DEPS-1 condensates are highly sensitive to mutator foci integrity. Functionally, *deps-1* is required for the steady state levels of secondary endo-siRNAs of piRNA and additional endo-siRNA pathways. Finally, we show that *deps-1* regulates small RNAs targeting P granule genes, suggesting small RNAs play a role in the maintenance of P granule equilibrium. Our study reveals that small RNA pathways and membraneless organelles are interdependent beyond the simple localization of small RNA components. Instead, an essential P granule factor actively participates in small RNA regulation by directly binding to an Argonaute protein.

## Results

### DEPS-1 and PRG-1 form intertwined ultrastructures

To identify proteins that intersect biomolecular condensate functions and small RNA pathways we performed immunoprecipitation (IP) of PRG-1 followed by liquid chromatography coupled with tandem mass spectrometry (LC-MS/MS, Figure S1A). This led us to identify 133 proteins to be preferentially in a complex with PRG-1 (Table S1). Of the 133 putative PRG-1 interactors, the P granule factor DEPS-1 (Defective P granules and Sterile-1) was among the ten most enriched factors (Figure S1B). DEPS-1 is a constitutive member of P granules and required for correct assembly of P granules (Spike et al., 2008) as well as for transgenerational inheritance (TEI) of exogenous RNAi (Houri-Ze’evi et al., 2016; Wan et al., 2018). First, we confirmed that PRG-1 and DEPS-1 colocalise in P granules by co-immunostaining transgenic animals expressing GFP-DEPS-1 (Figure 1A; Paix et al., 2014). In the adult germline, both proteins colocalise to P granules from the mitotic zone to the pachytene region. In the distal loop region where oogenesis begins and P granules start to disperse from the nuclear membrane (Updike and Strome, 2010) a higher portion of GFP-DEPS-1 starts to dissociate from the perinuclear region than PRG-1, suggesting the proteins are differentially regulated during a small temporal window. Given that PRG-1 binds to piRNAs to trigger secondary endo-siRNA biogenesis, we asked how and if the PRG-1/DEPS-1 complex interact with the mutator foci, a distinct biomolecular condensate that houses essential endo-siRNA components. While at lower resolution PRG-1 as well as DEPS-1 appear to be condensates that overlap with each other, these condensates can be further resolved to clusters of ultrastructures at higher resolution (Figure 1B). Often these DEPS-1 and PRG-1 ultrastructures weave around each other to form elongated condensates. In contrast, MUT-16 condensates do not resolve to smaller clusters and only juxtaposed close to the DEPS-1/PRG-1 complex (Figure 1B), consistent with mutator foci being an independent module (Phillips et al., 2012).

**Figure 1.**
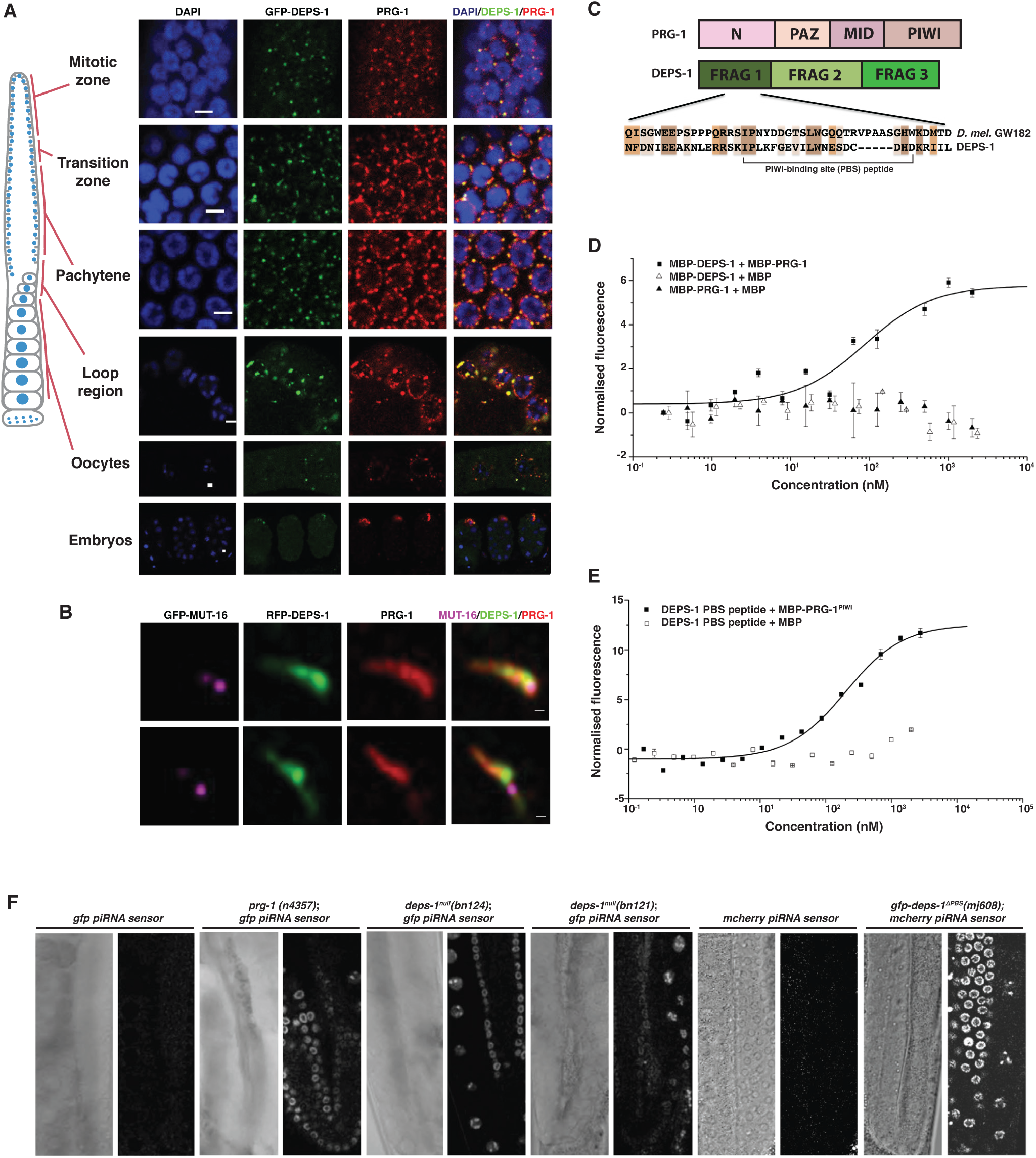
DEPS-1 is a direct interactor of PRG-1 and is required for efficient piRNA-mediated transgene silencing. A) DEPS-1 and PRG-1 colocalise as peri-nuclear granules. *C. elegans* germlines expressing GFP-DEPS-1 were dissected and immunostained for GFP and PRG-1. PRG-1 and DEPS-1 colocalise in the proliferative zone, transition zone, pachytene, oocytes and embryos. A higher proportion of PRG-1 remains perinuclear in the loop region compared to DEPS-1. Scale bar = 3 µm. B) Colocalisation of PRG-1, DEPS-1 and MUT-16. Dissected germlines co-stained for PRG-1, RFP-DEPS-1 and GFP-MUT-16 were imaged and deconvoluted with HyVolution settings. Two clusters with all three proteins present are shown. Scale bar = 0.25 µm. C) Domain/fragment architecture of PRG-1 and DEPS-1. Sequence alignment (Clustal W) of Ago binding motif II on *Drosophila melanogaster* GW182 and DEPS-1 PBS with flanking sequences. D) Recombinant full-length MBP-tagged PRG-1 and MBP-tagged DEPS-1 were purified for MST assays. A serial dilution of unlabelled MBP-PRG-1 was incubated with 10 nM of label MBP-DEPS-1 and tested for binding. The K_d_^app^ is 855+/−133 nM. E) MST measurement of fluorescently labelled DEPS-1 PBS motif peptide was incubated with unlabelled MBP-tagged PRG-1 PIWI domain. K_d_^app^ is 1.9 µM +/−98 nM. F) Mutations in *deps-1* lead to piRNA sensor transgene desilencing. Whitefield (right panel) and fluorescent (left panel) images of whole mounted animals show the piRNA sensor is efficiently silenced in wild-type animals and is desilenced in *prg-1 (n4357), deps-1* null (bn121 and bn124) and *deps-1*^Δ*PBS*^*::rfp* (mj650) mutants.

### DEPS-1 binds to PRG-1 via its Piwi Binding Site (PBS)

To investigate if the DEPS-1/PRG-1 interaction is direct and RNA-independent, we purified recombinant RNA-free full-length DEPS-1 and PRG-1 as MBP-fusion proteins and tested for binding using microscale thermophoresis (MST). DEPS-1 and PRG-1 interact with K_d_^app^ at 855 +/− 133 nM (Figure 1C and D; Figure S2A). So far only a few examples of direct interactors for the Ago protein family, in particular the PIWI clade, have been identified. A sub-micromolar dissociation constant is on par with GW182 and Tudor domain-containing proteins interaction with human Ago and mouse PIWI (Elkayam et al., 2017; Zhang et al., 2017) indicating DEPS-1 and PRG-1 binding is physiologically relevant.

To dissect the nature of the interaction, we truncated PRG-1 to its individual domains (Figure 1C), we identified the PIWI domain (PRG-1^PIWI^) to be responsible for binding to full-length DEPS-1 (Figures S2B and D). We were unable to rule out DEPS-1 interacting with the MID domain due to MBP-tagged MID domain of PRG-1 binds to the MBP tag non-specifically (Figure S2C). PRG-1^PIWI^ binds to full length DEPS-1 with K_d_^app^ = 349+/−45 nM which is comparable to the interactions with full-length PRG-1 suggesting that PRG-1 interacts with DEPS-1 with its PIWI domain.

DEPS-1 is not predicted to contain any known domain folds but consists of a poly-serine C-terminal end. Prediction for secondary structures suggests the protein is composed mainly of beta-sheets and loops (Figure S1D). Although many proteins required for P granule formation contain domains with long stretches of low complexity in amino acid composition (Banani et al., 2017), DEPS-1 is not predicted to possess such domains despite of its requirement for P granule integrity (Figure S1E). To identify which region of DEPS-1 is required for binding to PRG-1, we truncated DEPS-1 into three fragments of similar sizes and with fragment boundaries in regions lacking predicted secondary structures (Figure 1C). We detected binding between PRG-1^PIWI^ and the N-terminal fragment of DEPS-1 (DEPS-1^frag1^) only with a K_d_^app^ of 151+/−28 nM (Figures S2E and F). This is again in agreement with the binding between the full-length proteins indicating that DEPS-1 interacts with its N-terminal region with PRG-1 PIWI domain.

Given that DEPS-1 interacts with PRG-1 via PRG-1^PIWI^ and that the PIWI domains of PIWI and Ago families share similar folds overall (Matsumoto et al., 2016), we next asked if DEPS-1 shares any characteristics with known protein interactors of the Ago PIWI domain. The GW182 proteins have been shown to bind to Ago PIWI domain by multiple GW motifs which fit into two tryptophan-binding pockets (Takimoto et al., 2009; Pfaff et al., 2013; Sheu-Gruttadauria and MacRae, 2018). While DEPS-1 lacks any GW motifs, we noticed a degree of similarity between two short stretches of DEPS-1 and the *D. melanogaster* GW182 in our alignment (dmGW182), one at the N-terminal and the other the C-terminal of DEPS-1. The C-terminal region with similarity to dmGW182 is the poly-serine tail. Intriguingly, the N-terminal region of DEPS-1 similar to dmGW182 shares similarity with dmGW192’s Ago-binding motif II (Figure 1C; Elkayam et al., 2017; Eulalio et al., 2009). Moreover, this N-terminal region is contained within DEPS-1^frag1^ which binds to PRG-1^PIWI^. We therefore generated a peptide for part of this sequence (DEPS-1^peptide^; Figure 1C) to test its binding with PRG-1^PIWI^. DEPS-1^peptide^ binds to PRG-1^PIWI^ with a K_d_^app^ of 1.9+/−0.1 *µ*M indicating this Ago-binding motif II like region of DEPS-1 is indeed responsible for PRG-1 interaction (Figure 1E). We have therefore termed this motif as the Piwi-binding site (PBS).

### DEPS-1 is essential for piRNA function

Using the piRNA sensor, we next asked if *deps-1* functions in the piRNA pathway *in vivo*. The piRNA sensor is a genetic tool consisting of a GFP- or RFP-tagged histone 2B (H2B) with a piRNA target site at its 3’ end, rendering its expression dependent on the piRNA pathway (Bagijn et al., 2013). We analysed the effect of the PRG-1/DEPS-1 binding by removing the PBS from endogenous *deps-1* and replacing it with a 5x glycine residue-linker via Crispr-Cas9 gene editing (henceforth referred to as *deps-1*^Δ*PBS*^) as well as two *deps-1* null alleles (bn121 and bn124) (Spike et al., 2008). Crossing *deps-*1 mutants with *piRNA sensor* animals, we found that the piRNA sensor is de-silenced in both *deps-1* null alleles, as well as the *deps-1*^Δ*PBS*^ mutant, as in the *prg-1 (n4357)* mutant, indicating that *deps-1* and specifically its PBS is required for piRNA functions (Figure 1F). Correspondingly, small RNAs targeting the piRNA sensor are reduced in *deps-1*^*null*^ mutants (Figure S2G). Hence, DEPS-1 binding to PRG-1 is required for normal piRNA pathway activity.

### Uncoupling PRG-1 and DEPS-1 disrupts PRG-1 ultrastructure organisation

Having identified the PRG-1 binding site on DEPS-1 and shown that it is required for piRNA RNA functions, we asked how the removal of this site affects DEPS-1 localisation. Live imaging of GFP-DEPS-1^ΔPBS^ expressing animals revealed that in the absence of PBS, DEPS-1 becomes diffused in the cytoplasm and forms fewer granules. (Figure 2A). We imaged the condensates at a high resolution to inspect how the DEPS-1 and PRG-1 ultrastructures are affected. While DEPS-1^ΔPBS^ localises to PRG-1 condensates when DEPS-1 is able to associate with the peri-nuclear region, PRG-1 condensates contain either very little or no DEPS-1^ΔPBS^. Furthermore, PRG-1 ultrastructure does not intertwine DEPS-1^ΔPBS^ as it does with DEPS-1^WT^ ultrastructure. We measured the length of the PRG-1 condensates along the peri-nuclear edge and found that PRG-1 condensates (with and without DEPS-1^ΔPBS^) become more compacted compared with the PRG-1/DEPS-1^WT^ elongated condensates (Figure2B). Hence, the peri-nuclear arrangement of PRG-1 ultrastructure is maintained by the direct interaction with DEPS-1.

**Figure 2.**
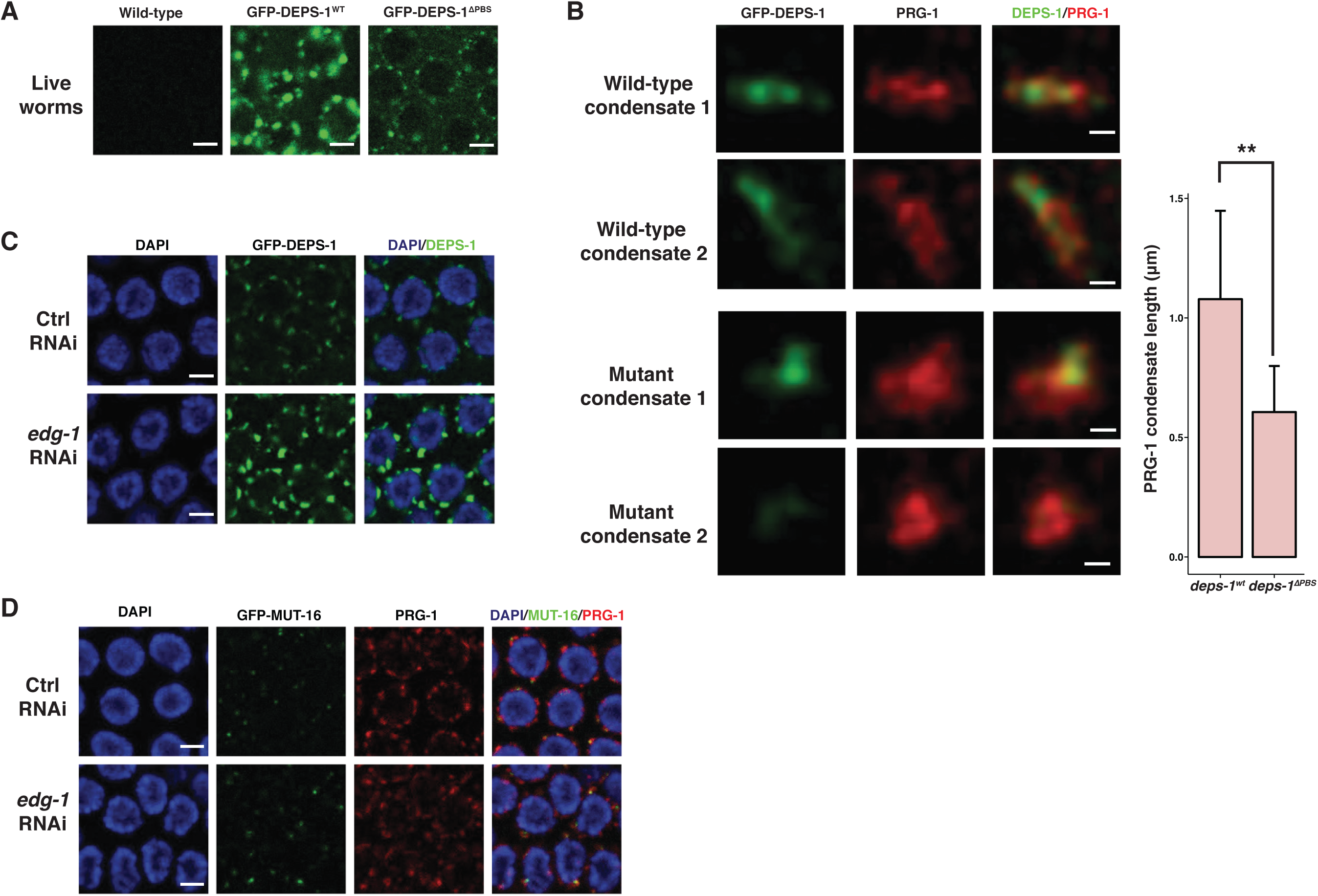
DEPS-1 condensates malform upon removal of PBS. A) Live worm imaging of the germline of wild-type animals (N2) (left), animals expressing wild-type GFP-DEPS-1 (*ax2063*; GFP-DEPS-1^WT^; middle) or GFP-DEPS-1 with mutated PBS (*mj608;* GFP-DEPS-1^ΔPBS^; right). GFP-DEPS-1^ΔPBS^ forms fewer granules and instead is diffused in the cytoplasm compared with GFP-DEPS-1^WT^. Scale bar = 3 µm. B) DEPS-1 and PRG-1 condensates are malformed in *gfp-deps-1*^Δ*PBS*^ *(mj608)* mutant. Dissected germlines co-stained for PRG-1 and GFP-DEPS-1 were imaged and deconvoluted with Hyvolution settings. Two selected clusters of each genotype were enlarged to show differences between the wild-type and PBS mutant form of GFP-DEPS-1. Top panels: *gfp-deps-1*^*WT*^ *(ax2063)*; Bottom panels: *gfp-deps-1*^Δ*PBS*^ *(mj608)*. The length of PRG-1 condensate along the perinuclear membrane was measured manually. Bar graph represents twenty PRG-1 condensates measured in 2 germlines (total n=40 for each genotype). Scale bar = 0.25 µm. C) *edg-1* was knocked down via RNAi in animals expressing *deps-1::gfp (ax2063)*. Animals were dissected for germline staining of GFP. DEPS-1 forms brighter granules upon *edg-1* knockdown. Scale bar = 3 µm. D) Animals expressing gfp::mut-16 (*mut-16 (pk710);mut-16::gfp*(*mgSi2))* were dissected and stained for GFP and PRG-1 after *edg-1* knockdown via RNAi. MUT-16 and PRG-1 granules were not affected by *edg-1* knockdown. Scale bar = 3 µm.

### EDG-1 interacts with both DEPS-1 and PRG-1 and modulates DEPS-1 condensates

Despite the uncoupling of DEPS-1 from PRG-1 by the deletion of PBS, DEPS-1^ΔPBS^ and PRG-1 remain in the same P granules at a low level. We therefore wondered if other proteins contribute to anchoring these proteins to the perinuclear region. We performed a yeast-two-hybrid (Y2H) screen using full length DEPS-1 as a bait. We obtained one high-confidence candidate, the putative protein encoded by *B0035.6*. Y2H data indicated that *B0035.6* interacts with DEPS-1 via its C-terminal region (Figure S3A). B0035.6 has no predicted conserved domain structures but a low similarity to human MEG-3 (Hubstenberger et al., 2015). B0035.6 was also one of the significant interactors of PRG-1 in our proteomic analysis (Table S1, Figure S3B), suggesting DEPS-1, B0035.6 and PRG-1 form a trimeric complex. We then performed RNAi knock-down of *B0035.6* via RNAi and tested if it is required for the normal formation of DEPS-1 or PRG-1 condensates. Reducing B0035.6 expression level lead to enlarged DEPS-1 condensates (Figure 2C), but not PRG-1 condensates (Figure 2D). We therefore named *B0035.6* as *<u>E</u>nlarged <u>D</u>eps <u>G</u>ranules-1 (edg-1)*. Given the dependence of the piRNA pathway on the mutator foci (Figure 2D), we tested if MUT-16 condensation was affected and found that they are not. Hence, while *edg-1* is found to be an interactor for both DEPS-1 and PRG-1, it specifically modulates DEPS-1 condensation.

### DEPS-1, PRG-1 and mutator condensates are interdependent

Mutations in P granule components were previously thought to be incapable of influencing mutator foci and vice versa (Phillips et al., 2012), even though materials such as RNAs must pass from P granules to mutator foci directly or indirectly as the piRNA pathway requires functional secondary endo-siRNA machineries to mediate gene silencing. Furthermore, Z-granules have recently been identified and shown to dynamically merge with P granules and mutator foci in different germline domains (Wan et al., 2018), indicating the physical separation between these distinct membraneless organelles is perhaps more fluid.

To understand the direct effect of DEPS-1 and PRG-1 interaction might contribute to the communication between P-granule and mutator foci, we investigated how DEPS-1, PRG-1 and MUT-16 condensates are affected by mutations in *deps-1, prg-1* and *mutator* genes. We noticed that the morphology of these perinuclear granules changes progressively in a transgenerational manner (Figure S4A), possibly related to the Mortal Germline defects observed in some piRNA pathway mutants (Billmyre et al., 2018), we therefore restricted our characterisation to the first five generations post-introduction of these mutations To carry out image analysis, we created an analytical pipeline consisting of a general-purpose object segmentation plugin (HKM Segment, https://github.com/gurdon-institute/HKM-Segment) for ImageJ (Rasband, 2014) called by a macro (https://github.com/gurdon-institute/HKM-Segment/blob/master/Kin_granules.ijm) to detect granules and measure intensity, area and circularity in our piRNA-condensate paradigm (Table 1).

**Table 1.** Summary of condensate defects in *prg-1, deps-1* and *mutator* mutants.

| Control strain genotype | Test strain genotype | Relevant mutation | Condensate | Condensate Defects |  |  |
| --- | --- | --- | --- | --- | --- | --- |
|  |  |  |  | Intensity | Area | Circularity |
| *deps-1::gfp (ax2063)* | *gfp::deps-1 (ax2063); mut-15 (tm1358)* | *mut-15* | GFP-DEPS-1 | ++<sup>down</sup> | +++<sup>down</sup> | ++<sup>up</sup> |
| *deps-1::gfp (ax2063)* | *gfp::deps-1 (ax2063); mut-16 (pk710)* | *mut-16* | GFP-DEPS-1 | +++<sup>down</sup> | +++<sup>down</sup> | +<sup>down</sup> |
| *deps-1::gfp (ax2063)* | *gfp::deps-1 (ax2063); mut-2(ne298)* | *mut-2* | GFP-DEPS-1 | +++<sup>down</sup> | +++<sup>down</sup> | - |
| *deps-1::gfp (ax2063)* | *gfp::deps-1 (ax2063); prg-1 (n4357)* | *prg-1* | GFP-DEPS-1 | +<sup>up</sup> | +++<sup>down</sup> | +++<sup>up</sup> |
| *deps-1::gfp (ax2063)* | *gfp::deps-1 (ax2063); mut-15 (tm1358)* | *mut-15* | PRG-1 | - | - | - |
| *deps-1::gfp (ax2063)* | *gfp::deps-1 (ax2063); mut-16 (pk710)* | *mut-16* | PRG-1 | - | +<sup>down</sup> | - |
| *deps-1::gfp (ax2063)* | *gfp::deps-1 (ax2063); mut-2(ne298)* | *mut-2* | PRG-1 | - | - | - |
| *mut-16 (pk710); mut-16::gfp (mgSi2 IV)* | *mut-16 (pk710); mut-16::gfp (mgSi2 IV); deps-1 (bn121)* | *deps-1* | PRG-1 | ++<sup>up</sup> | - | - |
| *deps-1::rfp (ax2063); mut-16 (pk710); mut-16::gfp (mgSi2 IV)* | *deps-1ΔPBS::rfp (mj605); mut-16 (pk710); mut-16::gfp (mgSi2 IV)* | *deps-1<sup>ΔPBS</sup>* | PRG-1 | +<sup>up</sup> | +<sup>down</sup> | ++<sup>up</sup> |
| *mut-16 (pk710); mut-16::gfp (mgSi2 IV)* | *mut-16 (pk710); mut-16::gfp (mgSi2 IV); deps-1 (bn121)* | *deps-1* | GFP-MUT-16 | +<sup>up</sup> | - | - |
| *deps-1::rfp (ax2063); mut-16 (pk710); mut-16::gfp (mgSi2 IV)* | *deps-1ΔPBS::rfp (mj605); mut-16 (pk710); mut-16::gfp (mgSi2 IV)* | *deps-1<sup>ΔPBS</sup>* | GFP-MUT-16 | +<sup>up</sup> | - | - |
| *mut-16 (pk710); mut-16::gfp (mgSi2 IV)* | *mut-16 (pk710); mut-16::gfp (mgSi2 IV); prg-1 (n4357)* | *prg-1* | GFP-MUT-16 | ++<sup>down</sup> | - | +<sup>down</sup> |

Footnote: +++ = 0.001 > p-values; ++ = 0.01 > p-values > 0.001; + = 0.05 > p-values; - = p-values > 0.05. <sup>down</sup> = Decreased and <sup>up</sup> = Increased in values compared with corresponding wild-type animals.

*deps-1*^*null*^ and *deps-1*^Δ*PBS*^ mutations lead to PRG-1 condensates becoming brighter as reflected by higher condensate intensity, suggesting either more proteins are present in the condensates or PRG-1 becomes more densely packed (Figure 3A and D). Since *deps-1* mutations have been shown to alter the levels of the mRNA and proteins of P granule factors (Spike et al., 2008), we investigated if *deps-1* mutations also affects *prg-1*. No significant differences in either mRNA or protein products of *prg-1* in *deps-1* mutants were detected (Figures S4B and C). Hence, the effects of *deps-1* on PRG-1 condensate are solely in the subcellular distribution of the protein. Intriguingly, *deps-1* mutants contain fewer and brighter MUT-16 condensates despite DEPS-1 being a P granule component (Figure 3A and Figure S4D).

**Figure 3.**
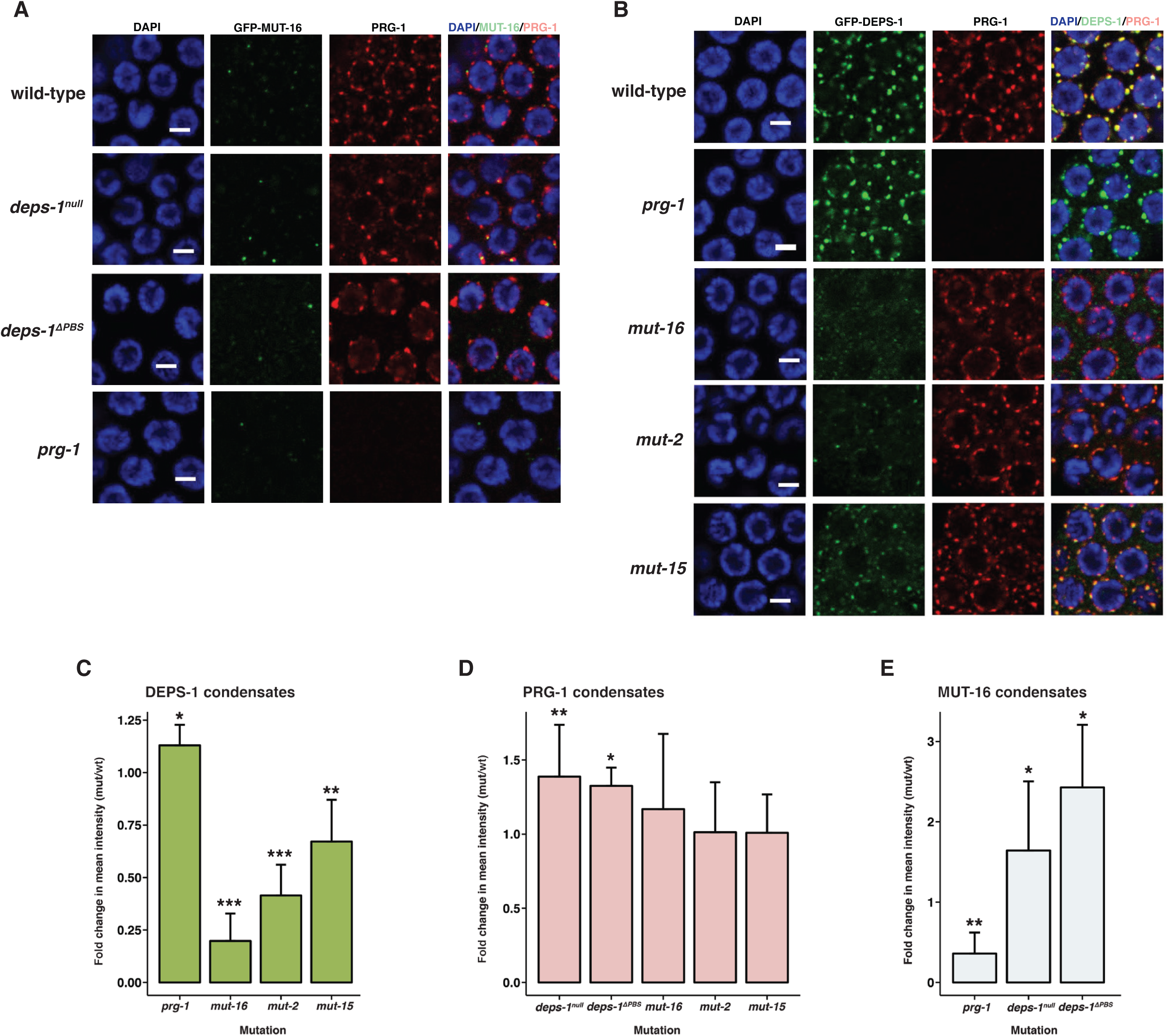
Morphologies of PRG-1, DEPS-1 and MUT-16 condensates are interdependent. A) GFP-MUT-16 and PRG-1 localisations were examined in *prg-1(n4357), deps-1*^*null*^*(bn121) and deps-1*^Δ*PBS*^*(mj605)* mutants. The common genotypes of strains used are *mut-16 (pk710); gfp::mut-16(mgSi2)* which is denoted as “wild-type”. Additional mutations upon this common genotype are indicated on the left. *C. elegans* germlines were dissected and immunostained for GFP and PRG-1. Scale bar = 3 µm. B) GFP-DEPS-1 and PRG-1 localisations were examined in *prg-1(n4357), mut-16(pk710), mut-2(ne298) and mut-15(tm1358)* mutants. “wild-type” indicates the common genotype of *gfp::deps-1(ax2063)* among the strains used and additional mutations are indicated on the left. *C. elegans* germlines were dissected and immunostained for GFP and PRG-1. Scale bar = 3 µm. C) – E) Graphs comparing DEPS-1 (C), PRG-1 (D) and MUT-16 (E) condensate intensity in various genotypes. Quantifications were obtained from 3 separate dissections of 2-4 germlines per dissection. Error bars represent standard deviations.* = 0.05> p-value >0.01; ** = 0.01> p-value >0.001; *** = 0.001> p-value.

Interruption of *prg-1* leads to mildly brighter peri-nuclear GFP-DEPS-1 condensates (Figure 3B). In contrast, removal of the PBS leads to DEPS-1^ΔPBS^ forming fewer condensates and becoming diffused in the cytoplasm (Figure 2A). This suggests that the PBS mediates the interaction of DEPS-1 with additional P granule components in addition to PRG-1 (see small RNA section). Dimmer MUT-16 condensates were found in *prg-1* mutant (Figure 3A, bottom panel), again even though PRG-1 resides in the P granules.

Mutations in either *mut-16* or *mut-2*, and to a lesser extent *mut-15*, abolish the perinuclear association of DEPS-1^WT^ (Figure 3B). Surprisingly, despite their effects on DEPS-1 position, *mut-16, mut-2* and *mut-15* mutations do not affect the intensity of PRG-1 condensates (Figure 3D). This indicates that the perinuclear localisation of PRG-1 is not dependent on the presence of DEPS-1 in the same condensate and is in agreement with the more intense PRG-1 condensate localised to the perinuclear region in *deps-1* mutants.

Interestingly, we found that in some instances where the brightness of the condensates are affected the circularity and area are also altered (Table1). These might reflect the rearrangements of ultrastructure within the condensate. Overall, we found that mutations of *deps-1* and *prg-1* affect MUT-16 condensate (Figure 3A and E and Table 1). Similarly, mutations in *mutator* genes lead to defects in DEPS-1 condensate (Figure 3B and C, Table 1). These observations indicate that piRNA pathway components located in P granules and mutator foci can be reciprocally or co-regulated.

### *deps-1* is required for the steady state levels of 22G RNAs against piRNA targets

To understand what role *deps-1* plays in the piRNA pathway, we profiled the small RNA populations in the *deps-1* mutants. We first examined if the 21Us are affected. Sequencing of the small RNA population from 5’-independent libraries show that a *deps-1 null* mutant has a comparable level of 21U population as in *wild-type* animals (Figure 4A). We then examined the 22G population in *deps-1* mutants. We find that the level of 22Gs against most *prg-1* targets is diminished (Figure 4B and Figure 5A) and that *deps-1* has a milder effect on 22Gs compared with *prg-1* and *mut-16* mutants, indicating that additional genes contribute to linking 21Us to 22Gs production. We conclude from the small RNAs characterisation of *deps-1* mutants that *deps-1* functions immediately downstream of *prg-1* and upstream of, or in parallel to secondary endo-siRNA production. Of note, DEPS-1 was not found as an interactor of HRDE-1 (Akay et al., 2017), an essential Ago protein that transports 22Gs into the nucleus to initiate nuclear gene silencing further suggesting that DEPS-1 does not function downstream of secondary endo-siRNA biogenesis.

**Figure 4.**
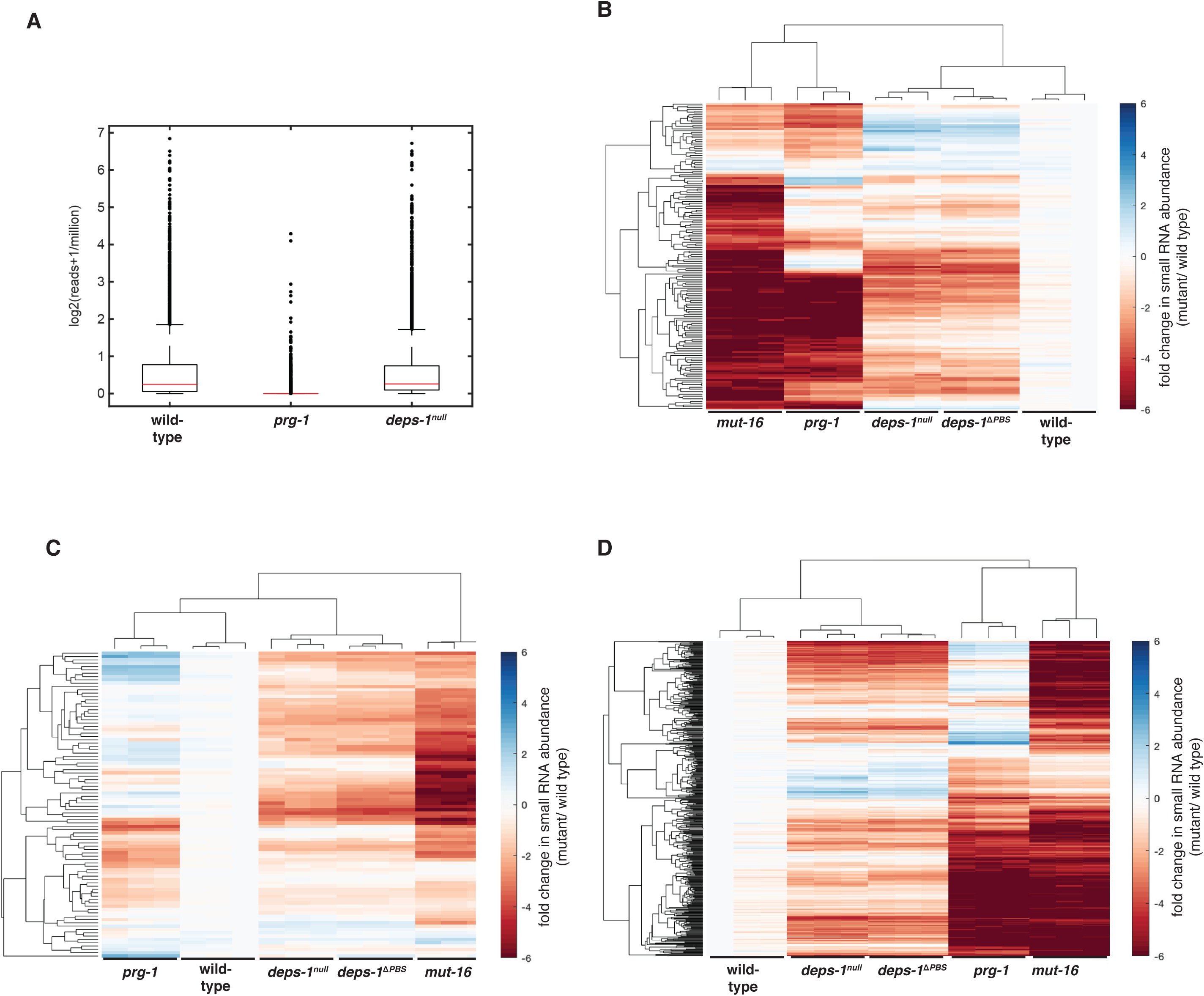
*deps-1* regulates 22Gs against piRNA targets and other endo-siRNAs. A) Small RNAs were sequenced in animals containing piRNA sensor (*mjIs144*; denoted as wild-type) alone, or in the presence of *deps-1(bn124)* or *prg-1(n4357)* mutations. *deps-1* mutant expresses similar level of 21Us as in wild-type animals, whereas 21Us in *prg-1(n4357)* mutant is significantly diminished compared with wild type and *deps-1* mutant (p-value <10^−20^). B) - D) Cluster analysis of 5’-independent small RNA libraries showing the fold change of small RNAs mapped to known targets of different small RNA pathways: piRNA targets (B), repetitive elements (C) and *wago* targets (D), in the indicated mutants compared to wild type. “wild-type” denotes animals expressing *piRNA sensor*(*mjIs144);* “*deps-1*^*null*^” denotes animals expressing *deps-1(bn121)*; *piRNA sensor* (*mjIs144);* “*deps-1*^Δ*PBS*^” denotes animals expressing *deps-1*^Δ*PBS*^ *(mj605); mut-16(pk710); mut-16::gfp(mgSi2);* “*mut-16*” denotes animals expressing *mut-16* (pk710); *piRNA sensor* (*mjIs144)*. Fold change is displayed in natural log.

**Figure 5.**
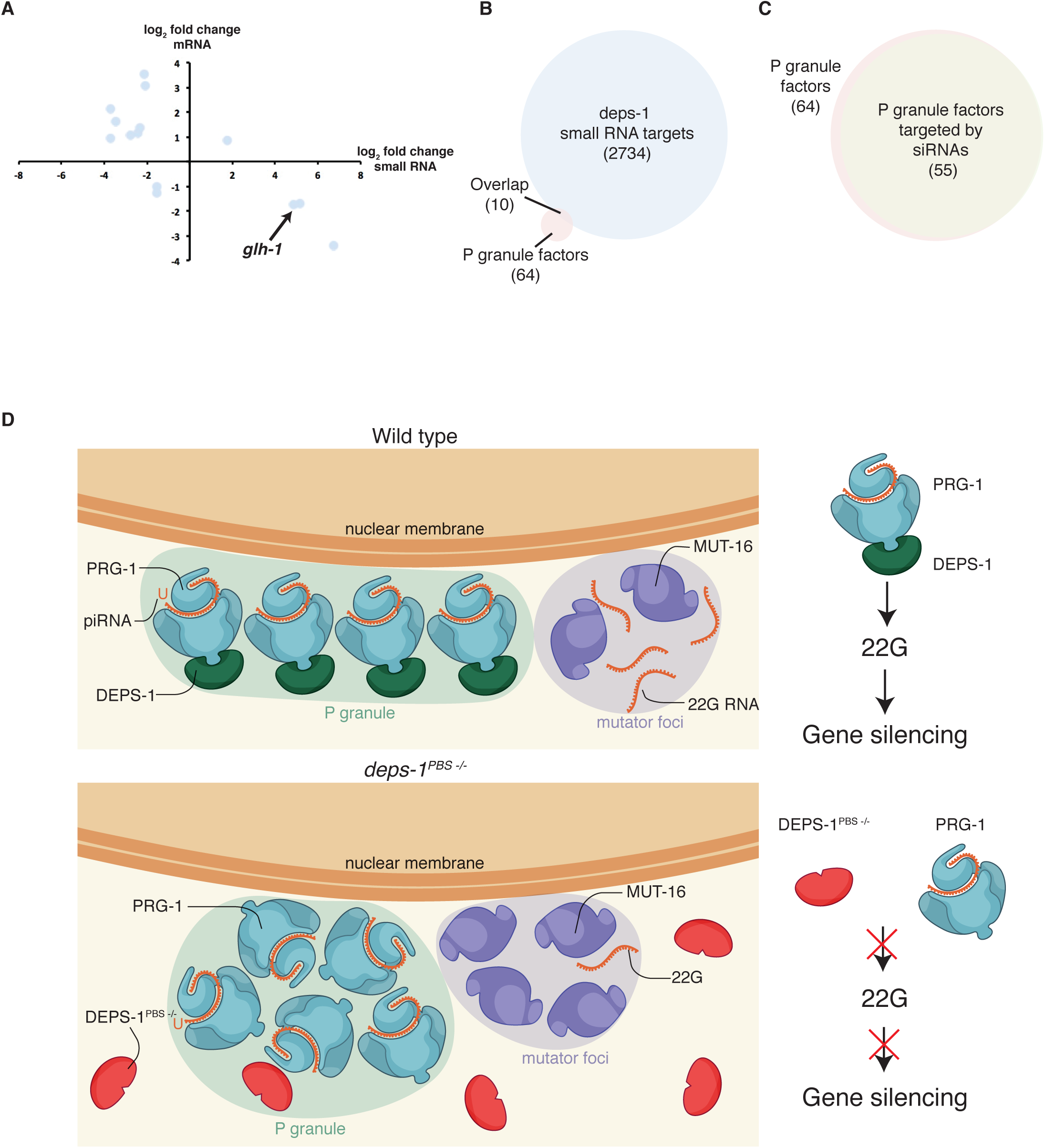
P granule components are targeted by small RNAs and mechanistic model of DEPS-1 function in small RNA pathways. A) Fold change in 22Gs and mRNA transcripts of differentially regulated genes in *deps-1* null mutant were compared. Fold changes of mRNA transcript were obtained from Spike et. al., 2008. Differential regulation of 22Gs is defined as adjusted p-value < 0.05, fold change compared to wild-type either <0.5 or >2. B) The 22Gs of 10 P granule factors are differentially regulated in *deps-1 (bn121)* mutant (p-Value < 0.001 Hypergeometric test). C) 58 P granule factors are targeted by endo-siRNAs (p-Value < 0.001 Hypergeometric test). D) Wild-type DEPS-1 binds directly to the PIWI domain of PRG-1 via the PBS motif in the N-terminal. The PRG-1 and DEPS-1 ultrastructures interact to form elongated perinuclear condensates. Intact PRG-1/DEPS-1 complex maintains normal morphology of PRG-1 and MUT-16 condensates. Efficient gene silencing mediated by the piRNA pathway ensues (top panel). DEPS-1 PBS mutant cannot associate with PRG-1 in perinuclear condensates and becomes dispersed in the cytoplasm. This results in an increase in the intensity of PRG-1 and MUT-16 condensates. Additionally, the steady-state level of secondary endo-siRNA from multiple pathways is reduced in the presence DEPS-1 PBS mutant. In particular, the piRNA pathway cannot mediate gene silencing (bottom panel).

### *deps-1* functions in multiple small RNA pathways

Since DEPS-1^ΔPBS^ localisation is significantly more disrupted than DEPS-1^WT^ in *prg-1* null mutant, we hypothesise that DEPS-1 interacts with other Ago proteins given that the PIWI domain is a conserved feature in the family of proteins and that multiple small RNA pathways function in the *C. elegans* germline (Grishok, 2013; Meister, 2013). To identify what other Ago proteins might cooperate with DEPS-1, we analysed if other small RNA populations are affected in *deps-1* mutants. We find that *deps-1* 22Gs profile targeting repetitive elements resembles that of the *mut-16* mutant in which 22Gs against repetitive elements that are independent of *prg-1* regulation are diminished (Figure 4C and Figure 5B). *deps-1* also affects subpopulations of *prg-1* independent *wago* targets (Figure 4D and Figure 5C; Gu et al., 2009). However, unlike *mut-16, deps-1* has limited effects on *ergo-1* targets which consist of recently duplicated genes (Figure 5D; Vasale et al., 2010). The effect of *deps-1* on small RNAs is not a general phenomenon due to its requirement in P granule maintenance, as it only mildly affects a small fraction of *csr-1* targets (Figure 5E; Conine et al., 2010), Therefore *deps-1* functions in multiple *mut-16*-dependent germline small RNA pathways.

Having seen that *deps-1*^Δ*PBS*^ has a similar effect on small RNAs as *deps-1* null mutation, we tested if *deps-1*^Δ*PBS*^ is also resistant to germline RNAi as observed in the *deps-1* null mutants (Spike et al., 2008). Knockdown of *pos-1* results in dead embryos. Indeed, *deps-1*^Δ*PBS*^ is resistant to RNAi in the germline (Figure S5H). While in *dep-1* null and *deps-1*^Δ*PBS*^ in general exhibit similar small RNA profiles and germline RNAi insensitivity, whether or not all of these small RNA functions are dependent on the direct interaction of *deps-1* with the responsible Ago proteins requires further biochemical characterisations.

### Small RNAs target P granule components as part of an auto-regulatory loop

It has previously been shown that the mRNAs of 13 and 32 genes are significantly downregulated and upregulated with the *deps-1* mutant, respectively (Spike et al., 2008). We therefore asked if the effect of *deps-1* on these genes could be mediated by small RNAs. We identified small RNAs targeting 14 of these genes as differentially regulated in our *deps-1* null mutant (Figure 5A). We found in general when small RNAs are downregulated, the mRNAs are upregulated in *deps-1* null mutant and vice versa, suggesting these effects are mediated by the perturbations in small RNA populations. One of the genes modulated as such is *glh-1*. GLH-1 is an essential P granule factor and its protein level is greatly reduced in *deps-1* mutant (Spike et al., 2008). Therefore we wondered if *deps-1* might regulate other P granule factors by modulating small RNA levels. We obtained a list of P granule factors from AmiGO under the GO term “P granule” and manually curated the list to remove protein isoforms of the same gene. This resulted in 64 factors identified as P granule components (Table S2). We found 2734 genes to be differentially targeted by *deps-1* (padj <0.05 with fold change below 0.5 or above 2 compared to *wild-type)* of which 10 are P granule factors (Figure 5B; Table S2). Strikingly, 55 out of 64 P granule factors are in fact endo-siRNA targets (Figure 5C; Table S2) suggesting P granules in addition to housing various small RNA pathways are themselves targets of these pathways potentially forming feedback loops.

## Discussion

Small RNA pathways associate with membraneless organelles to mediate their functions. Here we report for the first time the molecular details of how the piRNA pathway engages with the P granule by the direct coupling of the PIWI protein PRG-1 to the essential P granule component DEPS-1. We demonstrate that DEPS-1 interacts via a novel PIWI binding site (PBS) with PRG-1, which is required to maintain an elongated arrangement of PRG-1 ultrastructures. *deps-1* is required for subpopulations of secondary endo-siRNAs concomitant with a disruption of the mutator foci whose mutations reciprocally affect DEPS-1 protein distribution. Our study presents novel insights into how PIWI proteins function, as well as the intricate links between proteins and protein-nucleic acid interactions in complex liquid-liquid phase-separation organelles.

### Dynamics of P granules

The composition of *C. elegans* P granules is dynamic. For example, CAR-1-containing germline mRNA processing bodies (grPBs) associate with P granules only in oocytes (Hubstenberger et al., 2013); Wan et al. recently identified ZNFX-1 as a novel endo-siRNA factor that is spatially and temporally regulated to form Z-granules that fuse and dissociate from P granules and mutator foci (Wan et al., 2018). Although both are considered constitutive P granule components, we observed here that DEPS-1 is spatially regulated distinctly from PRG-1 during a small temporal window when oocytes are formed and P granules begin to disperse from the nuclear envelope. Whether or not this difference in spatial regulation is translated into small RNA regulation, given that *deps-1* regulates small RNAs independent of *prg-1* and vice versa, remains to be investigated. Furthermore, *edg-1* was previously identified as a modifier for grPBs and hence it possibly links grPBs to P granules (Hubstenberger et al., 2015). These highlight the dynamic nature of the proteins that constitute the P granules and also that distinct granules cooperate. It would therefore be interesting to understand how these dynamics assist small RNA biology and germline development.

### DEPS-1 and PRG-1 ultrastructure

Perinuclear germ granules are conserved features throughout the animal kingdom, acting as platforms for RNA metabolism and RNA-mediated gene regulation and are essential for germ cell development (Updike et al., 2011; Voronina et al., 2011). For example, depletion of P granules in *C. elegans* leads to the formation of neurite-like projections in germ cells indicating P granules maintain germ cell identity (Updike et al., 2014). Their liquid-like property is thought to facilitate dynamic internal rearrangements as well as exchange of materials with their surroundings. However, recently a non-dynamic, gel-like scaffold has been found to envelope the liquid-core of P granules (Putnam et al., 2019). Moreover, differences in translational activity between the periphery and the core of the P-bodies have been observed in *Drosophila* oocytes (Weil et al., 2012). These indicate substructures exist within membraneless organelles to support their functional complexities, just as their membrane-bound counterparts. We observe here that PRG-1 and DEPS-1 condensates form smaller substructures that intertwine to form elongated complexes. This conformation is dependent on the direct interaction of PRG-1 with DEPS-1 as PRG-1 condensates collapse into a more compact arrangement in *deps-1*^Δ*PBS*^ mutants. Whether and how the elongated shape of the DEPS-1/PRG-1 complex enhance its small RNA function is unclear. Given that the piRNA-loaded PRG-1 scans for target mRNAs emerging from the nucleus through the nuclear pores (Pitt et al., 2000; Updike et al., 2011), we speculate its elongated morphology increases the ability of the DEPS-1/PRG-1 complex to cover the nuclear pores.

### DEPS-1 PBS mediates PRG-1 binding

The Ago and PIWI clades of the Argonaute protein family are highly conserved throughout the animal kingdom. The structure of the bombyx PIWI protein confirms that piRNAs associate with PIWI in the same orientation as in miRNAs with the Ago clade proteins (Elkayam et al., 2012; Schirle et al., 2014; Matsumoto et al., 2016). Additionally, Shen et al. discovered that the laws governing the base-pairing between piRNAs and target mRNAs are similar to their miRNAs counterparts (Bartel, 2018; Shen et al., 2018). These suggest that the Argonaute protein family are not only similar in protein structure, the manner in which these proteins carry out their downstream functions are also similar. Given that Ago binds to GW182 via its PIWI domain to recruit deadenylase complexes (Jonas and Izaurralde, 2015), it is perhaps anticipated that DEPS-1 acting immediately downstream of PRG-1 would bind directly to PRG-1^PIWI^ via a motif similar to the Ago-hook on GW proteins. The fact that the PIWI domains in both Ago and PIWI proteins are used to engage with downstream processes suggests that protein orientation with respect to the 5’ region of the small RNAs (which is anchored to the MID-PIWI domain) is particularly important for downstream protein recruitment. Despite of its similarity with the Drosophila GW182 protein, DEPS-1 is not an ortholog of the GW protein family. It would thus be helpful to determine structurally which residues on PRG-1^PIWI^ are required for DEPS-1 binding.

### Role of DEPS-1 in P granule and mutator foci communication

Despite of the aberrant PRG-1 condensates in the *deps-1* null mutant, the piRNAs population is normal in these animals suggesting that 21Us are still able to associate with PRG-1 as the absence of PRG-1 abolishes piRNAs (Batista et al., 2008; Das et al., 2008). In contrast, *deps-1* mutations reduce endo-siRNAs of piRNAs origin and lead to brighter and fewer MUT-16 foci. Furthermore, despite the reduction in the number of PRG-1 and MUT-16 condensates in *deps-1* mutants, these condensates always juxtapose each other, suggesting MUT-16 condensates require materials from the PRG-1 condensates to nucleate. Therefore, we speculate that normal morphology of PRG-1 condensates is required to communicate with the endo-siRNA pathway components in the mutator foci and that abnormal MUT-16 condensate is the consequence of the disrupted flow of RNAs from P granules. While much focus has been placed on how proteins drive phase-transition in RNP foci formation, a flurry of recent studies investigated the importance of RNAs in the formation of RNP foci. Langdon et al. shows that the secondary structure of mRNAs plays essential roles in specifying distinct Whi3-containing RNPs (Langdon et al., 2018). Furthermore, RNA:protein ratios determine phase-transition events in proteins prone to solid aggregation (Maharana et al., 2018). Given that a myriad of small RNA pathways are routed through the P granules and mutator foci in *C. elegans* (Grishok, 2013), it will be interesting to decipher how small RNAs and their target mRNAs affect the normal function of these organelles by modulating their formation.

### Small RNA pathway functions of DEPS-1

The dependence on *deps-1* for the normal level of 22Gs, but not 21Us, against piRNA targets places *deps-1* downstream of *prg-1. deps-1* also plays a role in 22Gs targeting repetitive elements as well as certain *wago* targets. Incidentally, DEPS-1 localisation is more severely affected in *deps-1*^Δ*PBS*^ than in *prg-1* null mutants, together suggesting the PBS on DEPS-1 mediates interaction with other Ago proteins. Hence, DEPS-1 serves as a common factor in multiple small RNA pathways. Previous work shows that the absence of piRNAs leads to secondary endo-siRNAs being mis-loaded to Argonaute proteins (Phillips et al., 2015) and that the zinc-finger protein ZNFX-1 associates with multiple Argonaute proteins (Ishidate et al., 2018). Thus, balanced inputs of small RNAs compete for the small RNA machineries to achieve balanced epigenetic signals. DEPS-1 is therefore likely to be critical in the fine-tuning of piRNA and endo-siRNA pathways.

### P granule components regulated by small RNAs to maintain P granule equilibrium

Hubstengerger et al. comprehensively catalogued the proteins and RNAs contents of P-bodies and found that the mRNAs of P-bodies proteins are themselves enriched in P-bodies. This highlights a self-regulatory aspect of organelles whose formation are highly dependent on concentrations of their constituents (Hubstenberger et al., 2017). Similarly, we found in our analysis that most P granule factors in *C. elegans* are siRNA targets and no single small RNA pathway is responsible as collectively they are targeted by multiple Ago proteins. This strategy likely prevents the malfunction of one small RNA pathway from causing the collapse of all P granule functions.

## Acknowledgements

We are grateful for the generous gifts of the DEPS-1::GFP and DEPS-1::RFP - expressing strains from the Seydoux Laboratory and the Mutator strains from the Gary Ruvkun laboratory. RNAi-expression clones were kind gifts from the Ahringer laboratory. We are indebted to Dr. Nicola Lawrence from The Gurdon Institute Imaging Facility for her help in high-resolution imaging. Some strains were provided by the CGC, which is funded by NIH Office of Research Infrastructure Programs (P40 OD010440). We thank Kay Harnish of the Gurdon Institute Sequencing Facility for managing the high-throughput sequencing.

## Materials and Methods

### Immunoprecipitation for mass spectrometry

Synchronized wild-type N2 and *prg-1 (n4357)* animals were grown to 1 day-old adults at 20 °C on HB101. After washing thoroughly to remove bacteria, animals were resuspended in lysis buffer (20mM HEPES, pH7.5, 150mM NaCl, 0.5% NP-40) and snapped frozen in liquid nitrogen. The samples were then lysed by bead-beating, followed by centrifugation at 4 °C to remove insoluble debris. Anti-PRG-1 antibody (Custom) or rabbit IgG was pre-coupled to protein A/G matrix (Thermo Scientific, 88802) and incubated with the supernatant of worm lysates for 4 h (Anti-PRG-1 with N2 lysates, anti-PRG-1 with *prg-1(n4357)* lysates and anti-IgG with N2 lysates). The immunoprecipitants were then washed with 3×1 ml of lysis buffer and eluted in elution buffer (8 M urea, 10 mM HEPES pH 8.0) with shaking at room temperature for 30 mins.

### LS-MS/MS

Experiments were carried out as described before (Bensaddek et al., 2016). Briefly, proteins eluted from immunoprecipitations were reduced and alkylated. Quantified proteins were then digested consecutively in solution using Lys-C and trypsin (both at 1:50 enzyme:substrate ratio). Peptides were desalted, dried and redissovled in 5% formic acid. RP-LC was performed using a multistep gradient from 5% solvent B to 35% solvent B. Solvents A and B were 2% ACN with 0.1% FA and 80% ACN with 0.1% FA, respectively. MS/MS data were acquired on a Q-Exactive Orbitrap in a data dependent mode.

### Mass spectrometery data analysis

Raw MS data were processed by MaxQuant (Cox and Mann, 2008). iBAQ values were divided by the total sum of intensity of each sample (Schwanhüusser et al., 2011). These normalised values were then log_10_ transformed to obtain normality and the resulting values were used for student’s t-test. To identify proteins enriched in immunoprecipitated PRG-1 from wild-type animals, the median of the transformed values were used for fold-change calculations.

### Molecular cloning and recombinant protein expression

All PRG-1 and DEPS-1 constructs were cloned using restriction enzymes into the pMAL-C5X vector. Recombinant proteins were expressed in BL21 (DE3) at 37 °C. Briefly, 10 ml of overnight pre-cultures were inoculated to 1 L of LB. Cells were grown to OD_600_ ∼0.8 and 1 mM IPTG was added to induce protein expression for 4 h. To purify the proteins, bacterial cells were lysed by sonication in PBS supplemented with protease inhibitors. 20 µg/ml RNaseA was added to the cleared lysates and incubated with gentle rotation overnight. Lysates were then applied to amylose resins (NEB, E8021) and washed with 20 column volumes of binding buffer supplemented with 14 mM beta-mercaptoethanol. Proteins were eluted with binding buffer supplemented with 20 mM maltose and 14 mM beta-mercaptoethanol. Proteins eluted from affinity column were subjected to size-exclusion chromatography equilibrated in assay buffer (30 mM HEPES pH 7.5, 100 mM K-Ac, 2mM Mg-Ac, 14 mM BME). Proteins used for MST assays were >85% in purity.

### Microscale thermophoresis (MST)

Proteins/peptides were labelled with atto-488 NHS ester (Sigma, 41698). Free dye was separated from labelled protein using G25 desalting columns. Unlabelled proteins were serial diluted 1:2 and incubated with a constant amount of labelled protein. MST assays were carried in assay buffer supplemented with 0.01% NP-40. Fluorescence was monitored throughout the assay (5 s laser off, 30 s laser on, 5 s laser off). The apparent dissociation constant (K_d_^app^) was calculated by the law of mass action using data from either thermophoresis or thermophoresis with temperature jump.

### Small RNA library preparation

Synchronized animals were grown to 1 day-old adults 20 °C. After washing thoroughly to remove bacteria with M9, animals were resuspended in TRIsure (Bioline, BIO-38033). Animals were lysed with 5x freeze-thaw cycles in liquid nitrogen. Total RNA was isolated by chloroform extraction. For 5’-independent libraries, 5 µg of total RNA was treated with 5’ polyphosphatase (Epicenter, RP8092H). Small RNAs were indexed using the TruSeq small RNA sample kit (Illumina) and size selected by gel separation in 6% TBE gels (Life Tech) and subsequently purified.

### Small RNA analysis

Small RNA sequencing results were obtained from https://basespace.illumina.com/ as fastq files after demultiplexing. Sequencing data is available in the European Nucleotide Archive under study accession number PRJEB31348 (Table S4). 3’ Adapter, reads below 18 nt length and reads with a length above 32 were removed using cutadapt. Remaining reads were aligned using STAR against the *C. elegans* genome WS235 allowing multimapping reads. To detect piRNAs, reads of 5’ dependent libraries were mapped against piRNA annotation WS235. Next, piRNA reads were counted using featurecount and abundance was calculated by correcting for library size using unique mapping reads. To compare piRNA sensor read distribution, reads of 5’ independent libraries were mapped against the piRNA sensor using STAR. Small RNA abundance was calculated by correcting for library size using H2B mapping reads. To compare small RNA changes in between worm strains of different gene set, small RNA reads per gene of a specific gene set were counted and abundance calculated by correcting for library size using unique mapping reads (cutoff > 50 reads per million). The mean smallRNA abundance per gene was calculated, next the fold-change was calculated by divided the mean abundance in *mutant* animals by the mean abundance in wild-type animals. Gene sets were obtained from Bagijn et al. 2012 for piRNA targets (Bagijn et al., 2013), Gu et al. (2009) for soma, germline and wago (Gu et al., 2009), Claycomb et al. (2009) for csr-1, Conine et al. (2010) for alg-3/4 (Conine et al., 2010),Vasale et al. (2010) for ergo-1 (Vasale et al., 2010), Buckley et al. (2012) for hrde-1 (Buckley et al., 2012) and repetitive element genes were detected using RepeatMasker.

### Worm dissection and immunostaining

1 day-old adult animals were dissected for germline and freeze cracked on poly-lysine coated microscope slides. Dissected germlines were fixed in −20 °C methanol for 20 min. Fixed samples were washed with PBS-T (PBS supplemented with 1% tween-20) prior to primary antibody addition. Primary antibodies were incubated with the samples at 4 °C for overnight. Secondary antibodies were incubated at 37 °C for 1hr in the dark. Antibodies used: anti-PRG-1 (Custom, 1:1000); anti-mouse GFP (Thermofisher, A-11120; 1:400); OIC1D4 (Developmental Studies Hybridoma Bank; 1:50); anti-RFP. All fluorescence secondary antibodies were from Invitrogen and used at 1:500. Dissected and stained germlines were mounted with Vectorshield antifading agent supplemented with DAPI.

### Confocal microscopy

Images were taken on Leica SP8 confocal microscope. Images taken for granule quantification were of single slices, with pinhole set at 1 AU. HyVolution images were taken with pinhole narrowed to 0.5 AU to result in higher resolution. HyVolution images were deconvoluted using the HyVolution software.

### Granule pipeline for confocal image analysis

HKM Segment is a plugin for ImageJ inspired by Alexandre Dufour’s Hierarchical K-Means segmentation algorithm (Dufour et al., 2008) available in Icy (De Chaumont et al., 2012). In this implementation, agglomerative K-Means clustering is applied to the image histogram to determine K threshold levels. An initial set of *K*_0_intensity levels are initially assigned evenly spaced through the intensity range *r* = *i*_*max*_ *– i*_*min*_ regardless of frequency, and the merging distance 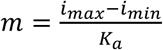 is recalculated for each iteration (a). Clustering is continued until assignment converges to give no further change in levels or cluster assignments, with *2* ≤ *K*_a_≤ *K*_0_. *K*_0_ is therefore the maximum permitted number of clusters and can be set as high as necessary to ensure separation of useful intensity classes, although increasing starting values will converge to the same final *K*_a_ when 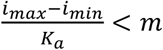.

The calculated threshold levels are applied in ascending order to extract objects within the specified size range. A thresholding algorithm can be chosen to filter out objects of low intensity, giving robust results in biological images without requiring subsequent level-sets segmentation as used by Dufour et. al. The fragments extracted by this method are further clustered to reconstruct objects that remain within the specified size range.

We used a custom ImageJ macro to call HKM Segment and output the results together with additional analysis. Regions of interest created by HKM Segment are output to the ImageJ Roi Manager when it is run from a macro, giving flexibility in the downstream analysis applied to segmented images. In this case, our macro measures the area, circularity and distance from the nearest nucleus boundary for granules detected with HKM Segment. The parameters used were a starting K of 16, a radius range of 0.1-0.6 µm and the Otsu thresholding method (Otsu, 1979).

Images were first manually curated to measure only areas containing germline, complete nuclei and nuclei at the widest cross section before measurements were performed using our macro.

Comparisons between different strains were usually obtained from three biological replicates, 2-4 germlines per replicate from 15 dissected germlines. Control and test strains were dissected and imaged in one setting. To calculate significant differences, mean values of each germline were obtained and tested for normality using the Shapiro-Wilk test. Student’s T-tests were performed on normally distributed data. Kolmogorov-Smirnov tests were performed on data not normally distributed. For fold change calculations, average intensity of all controls of one experiment was obtained and used to calculate the fold change of the individual mutant germline within the same experiment.

### Western blotting

Proteins from 75-150 µg of worm lysates were resolved by SDS-PAGE and transferred onto PVDF membrane. Antibodies used: Antibodies used: anti-PRG-1 (Custom, 1:1000; (Weick et al., 2014)); anti-tubulin (Sigma, DM1A; 1:1000).

### RNAi

Adult animals were bleached to obtain embryos which then hatched and synchronised in M9 for 24 - 48 h at 20 °C. L1 animals were fed with bacteria expressing control dsRNA or dsRNA against *edg-1*. 1-day old adults were subsequently dissected for germline imaging.

### General animal maintenance

Animals were fed with HB101 and maintained at 20 °C (unless stated otherwise) on NGM plates. Strains used in this study is listed in Table S4.

## Supplemental Figure Legends

**Figure S1.**
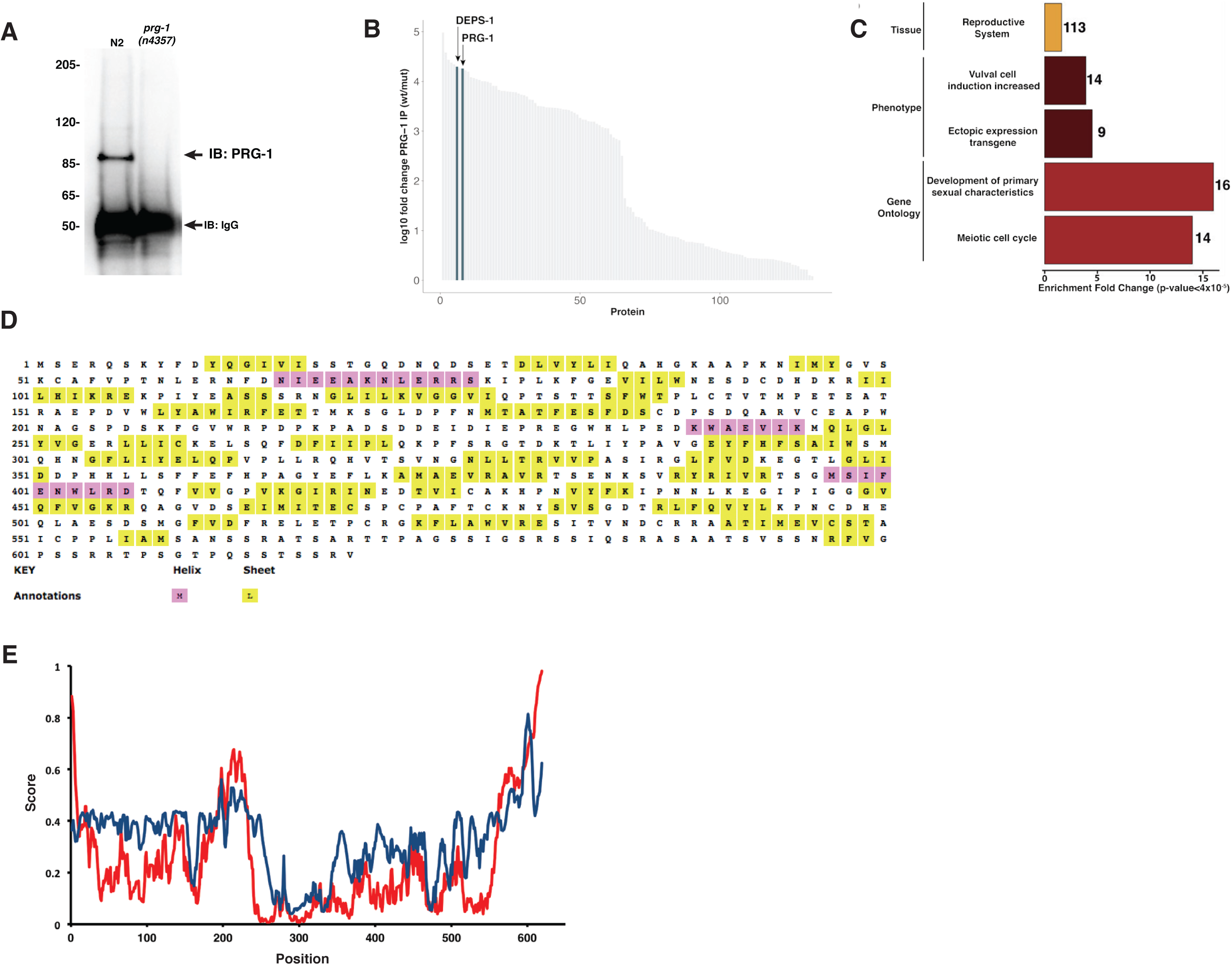
related to main figure 1. A) PRG-1 immunoprecipitation for mass spectrometry analysis. Anti-PRG-1 antibody immobilised on Protein A/G magnetic beads was incubated with lysates from wild-type or *prg-1(n4357)* animals. Western blot analysis of IPs shows PRG-1 was successfully pulldown. B) Proteins co-immunoprecipitated with endogenous PRG-1 were identified by LC-MS/MS. Median normalised iBAQ values were used to calculate fold-change enrichment in PRG-1 IP. Proteins enriched in PRG-1 IP with p-values <0.05 are ranked according to fold-change. B) Proteins enriched in PRG-IP with p-values<0.05 and enrichment fold change >1.5 compared to negative controls were analysed for tissue, phenotypic and gene ontology enrichment terms. Number at each bar indicates number of genes contribute to the enriched terms. D) and E) *In Silico* analysis of DEPS-1 structure. Secondary structure prediction of DEPS-1 by PSIPRED shows DEPS-1 is rich in β-sheets. Red: α-helix; Yellow: β-sheet. (D) Disorder prediction conducted by IUPRED suggests DEPS-1 does not contain large segments of disordered region (D).

**Figure S2.**
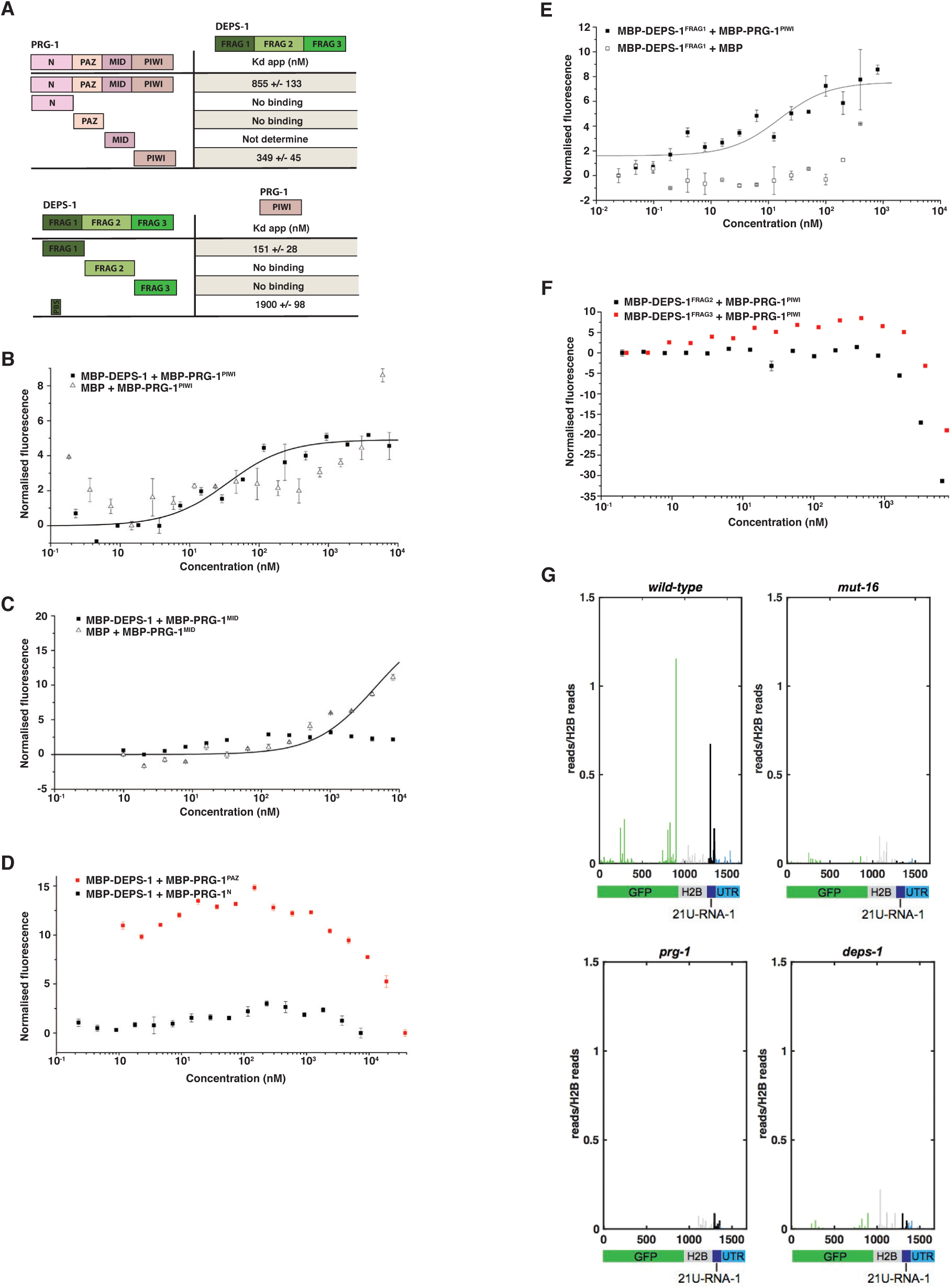
related to main figure 1. A) Summary table of K_d_^app^ of interactions between full length and truncated domains of DEPS-1 and PRG-1. B) - D) Full-length MBP-tagged DEPS-1 was fluorescently labelled and tested for binding with unlabeled MBP-tagged PRG-1^PIWI^ (B), PRG-1^MID^ (C), PRG-1^PAZ^ and PRG-1^N^ (D) using MST. The K_d_^app^ for binding between full length DEPS-1 and PRG-1PIWI is 349 +/−45 nM. E) and F) MBP-tagged PRG-1^PIWI^ domain was labelled and tested for interaction with unlabeled MBP-1 DEPS-1^frag1^ (E), DEPS-1^frag-2^ or DEPS-1^frag-3^ (F) using MST. The K_d_^app^ for binding between DEPS-1^frag1^ and PRG-1^PIWI^ is 151+/−28 nM. G) Analysis of small RNAs mapped to the piRNA sensor in wild type, *mut-16, prg-1* and *deps-1 mutants*

**Figure S3.**
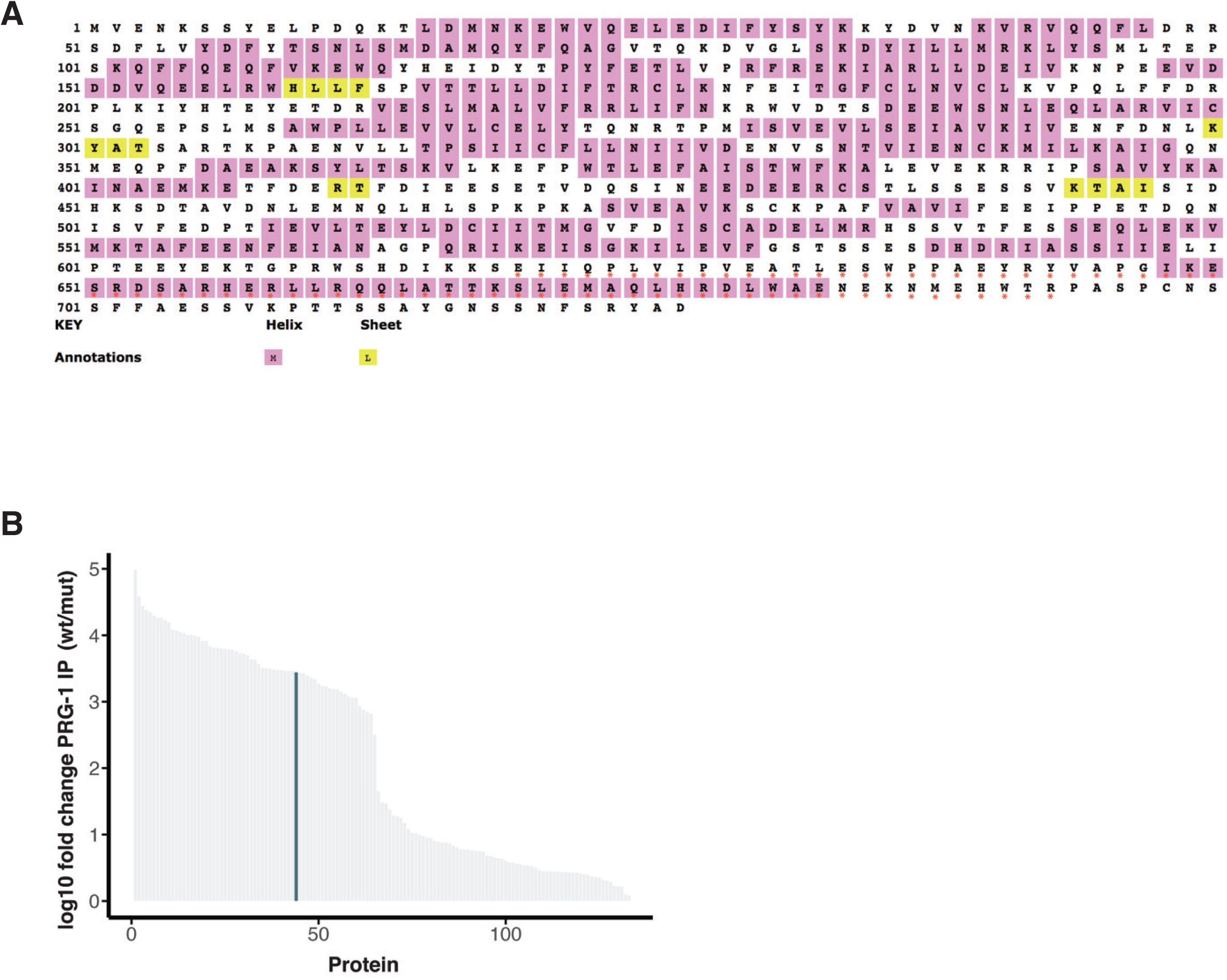
related to main figure 2. A) EDG-1 is predicted to be rich in α-helix by PSIPRED and lacks disordered region. Red asterisks highlight the minimal segment required for binding to DEPS-1 in yeast-two-hybrid screen. B) EDG-1 (highlighted in grey) was enriched in PRG-1 IP as identified by MS/MS.

**Figure S4.**
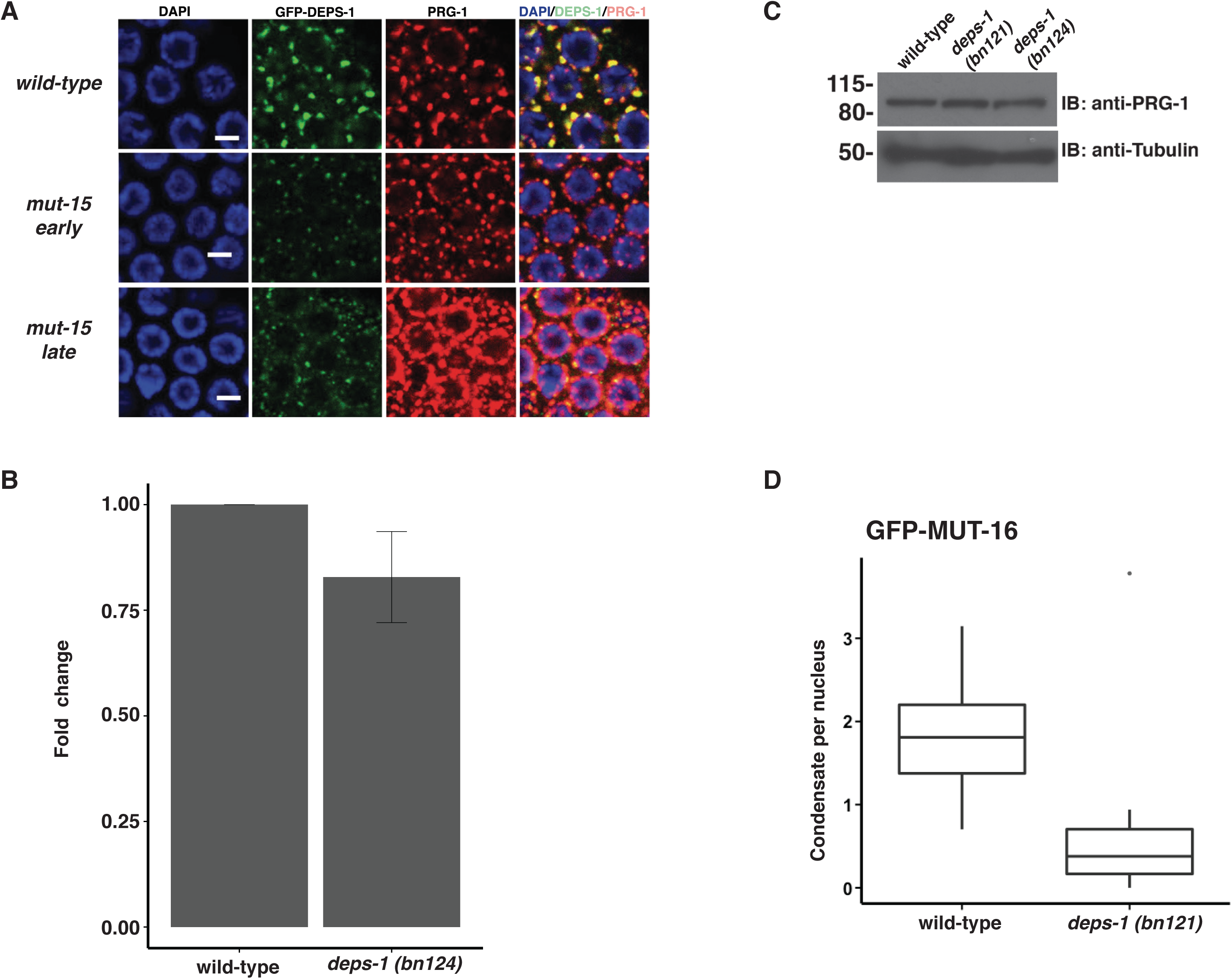
related to main figure 3. A) Defects in PRG-1 perinuclear accumulation in late generation of *mut-15* mutant. Dissected germlines co-stained for PRG-1 and GFP-DEPS-1 in wild-type and *mut-15*(*tm1358)* mutant animals. “wild-type” indicates the common genotype of *gfp::deps-1(ax2063)* among the strains used. Accumulation of PRG-1 to the perinuclear region in *mut-15* mutant is comparable to wild-type animals in early generations but is strongly affected in late generations. Scale bar = 3 µm. B) and C) PRG-1 mRNA and protein levels are not altered in *deps-1* mutant animal. RT-qPCRs were performed on N2 *wild-type* and *deps-1(bn124)* mutant animals. *prg-1* transcript levels were normalised to *act-3*. Average values of technical replicates of 3 of each biological replicate were used. Error bars are standard deviation of biological replicate of 2 (B). PRG-1 protein levels were assessed in N2 wild-type, *deps-1(bn121) and deps-1(bn124)* mutant animals by western blotting. Representative of 2 biological replicates (C).

**Figure S5.**
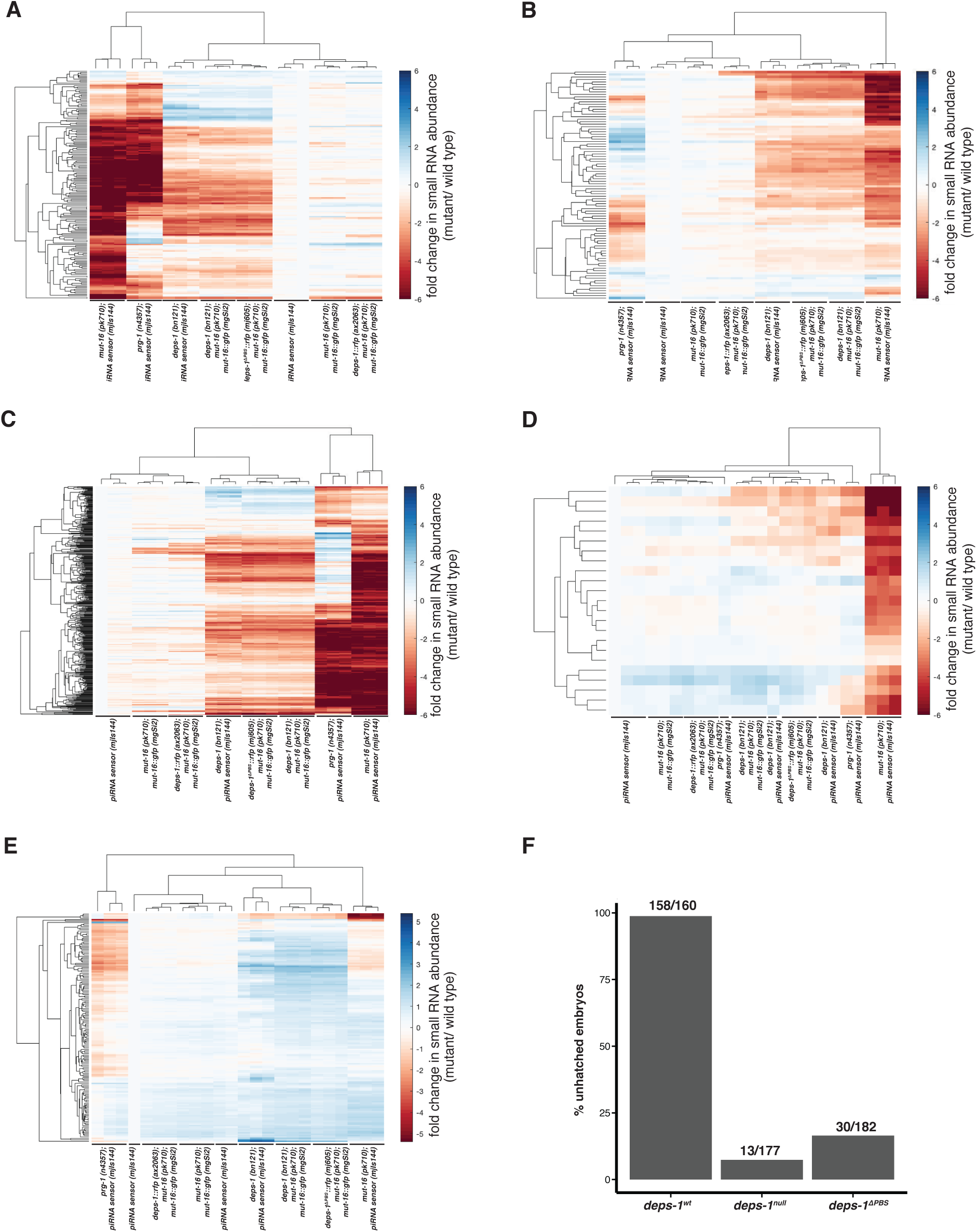
related to main figure 4. A) - G) Cluster analysis of small RNA libraries showing the fold change of small RNAs mapped to known targets of different small RNA pathways: piRNA targets (A), repetitive elements (B), wago12 targets (C), *ergo-1* targets (D), *csr-1* targets (E), in the indicated mutants compared to wild type. Fold change is displayed in natural log. H) *deps-1*^Δ*PBS*^ animals are germline RNAi incompetent. *pos-1* was targeted by RNAi in animals expressing either *rfp-deps-1*^*WT*^, *rfp-deps-1*^Δ*PBS*^ mutant or *deps-1*^*null*^*(bn121)* in the presence of *gfp-mut-16*; *mut-16(pk710)*. *rfp-deps-1*^Δ*PBS*^ mutant animals exhibited similar level of RNAi competency as *deps-1*^*null*^*(bn121)* while *rfp-deps-1* ^*WT*^ animals can fully mediate germline RNAi.

**Table S1 Putative PRG-1 interacting partners identified by LC-MS/MS.** Proteins identified with p-values<0.05 are listed.

**Table S2 List of P granule factors targeted by small RNA pathways.**

**Table S3 List of strains used in this study.**

**Table S4 Sample description of small RNA sequencing data.**

